# Bayesian Selection of Relaxed-clock Models: Distinguishing Between Independent and Autocorrelated Rates

**DOI:** 10.1101/2024.04.10.588547

**Authors:** Muthukumaran Panchaksaram, Lucas Freitas, Mario dos Reis

## Abstract

In Bayesian molecular-clock dating of species divergences, rate models are used to construct the prior on the molecular evolutionary rates for branches in the phylogeny, with independent and autocorrelated rate models being commonly used. The two classes of models, however, can result in markedly different divergence time estimates for the same dataset, and thus selecting the best rate model appears important for obtaining reliable inferences of divergence times. However, the properties of Bayesian rate model selection are not well understood, in particular when the number of sequence partitions analysed increases and when age calibrations (such as fossil calibrations) are misspecified. Furthermore, Bayesian rate model selection is computationally expensive as it requires calculation of marginal likelihoods by MCMC sampling, and therefore methods that can speed up the model selection procedure without compromising its accuracy are desirable. In this study, we use a combination of computer simulations and real data analysis to investigate the statistical behaviour of Bayesian rate model selection and we also explore approximations of the likelihood to improve computational efficiency in large phylogenomic datasets. Our simulations demonstrate that the posterior probability for the correct rate model converges to one as more molecular sequence partitions are analysed and when no calibrations are used, as expected due to asymptotic Bayesian model selection theory. Furthermore, we also show the model selection procedure is robust to slight misspecification of calibrations, and reliable inference of the correct rate model is possible in this case. However, we show that when calibrations are seriously misspecified, calculated model probabilities are completely wrong and may converge to one for the wrong rate model. Finally, we demonstrate that approximating the phylogenetic likelihood under an arcsine branch-length transform can dramatically reduce the computational cost of rate model selection without compromising accuracy. We test the approximate procedure on two large phylogenies of primates (372 species) and flowering plants (644 species), replicating results obtained on smaller datasets using exact likelihood. Our findings and methodology can assist users in selecting the optimal rate model for estimating times and rates along the Tree of Life.

## Introduction

Models of evolutionary rate variation among lineages, or clock models, are important because they are used to construct the rate prior in Bayesian inference of species divergence times (Thorne et al. 1998; Drummond et al. 2006; Rannala and Yang 2007). Roughly, three models (or classes of models) are commonly used: The strict clock, the independent-rates, and the autocorrelated-rates models. In the strict clock model, rates among branches in the phylogeny are equal (Zuckerkandl and Pauling 1965), an assumption that appears useful in the analysis of closely related species in which rates among branches are very similar (Yang and Rannala 2006). In the independent-rates model, rates among branches are independent, identical draws from a probability distribution such as the log-normal or the gamma distributions (Drummond et al. 2006; Rannala and Yang 2007), whereas in the autocorrelated-rates model, rates among branches are generated by an autocorrelated stochastic process such as the geometric Brownian motion process (Thorne et al. 1998; Rannala and Yang 2007) or the Cox-Ingersoll-Ross process (Lepage et al. 2007). A plethora of rate variation models have been developed encompassing the three classes of models or combinations of them (Huelsenbeck et al. 2000; Drummond and Suchard 2010; Li and Drummond 2012; Heath et al. 2012; Ho and Duchêne 2014; Lartillot et al. 2016; Fisher et al. 2023).

Choosing an appropriate rate model appears important because the different models can produce different divergence time estimates on the same dataset (e.g., dos Reis et al. 2018), and because different datasets may support different models. For example, Linder et al. (2011) found best support for the independent-rates model in flowering plants, but best support for the autocorrelated-rates model in Primates, when no fossil calibrations were used. Similarly, dos Reis et al. (2018) found a preference for the autocorrelated-rates model in Primates, while Barba-Montoya et al. (2018) found best support for the independent-rates model in flowering plants. Lepage et al. (2007) found autocorrelated rates had the better fit in eukaryotic, vertebrate and mammal datasets (see also McGowen et al. 2020; Álvarez-Carretero et al. 2022). Surprisingly, distinguishing between rate models was found to be difficult in simulated data (Ho et al. 2015), a puzzling result given that support for specific rate models has been strong in the real data analyses. The reasons for this discrepancy are not clear but at least two factors appear to limit our ability to distinguish between rate models: the non-identifiability of times and rates in Bayesian inference and sampling errors in the calculation of marginal likelihoods. Furthermore, real data analyses have been carried out on phylogenies with a few dozen taxa, and it is not clear whether support for a given rate model would still be maintained for larger phylogenies with hundreds of taxa. Analyses on large phylogenies have not been carried out because Bayesian model selection would be too computationally expensive on these datasets (e.g., Álvarez-Carretero et al. 2022), and thus inference technologies that can speed up analysis on large datasets are desirable.

Here we study the statistical properties of Bayesian selection of rate models for inference of divergence times. First, we use computer simulations to study the asymptotic properties of the model selection approach when the amount of data (the number of loci or alignment partitions) increases. Asymptotic analysis is important as it allows us to test whether a statistical procedure is reliable and accurate, that is, whether the procedure converges to the truth as the amount of data increases. Here we show that, when no fossil calibrations are used or when fossil calibrations are slightly mis-specified, Bayesian inference is well-behaved and Bayesian model selection asymptotically selects the correct rate model. On the other hand, when fossil calibrations are badly misspecified, the model selection approach is unreliable and the wrong model may be selected even with large datasets. We also study the use of likelihood approximations to speed up computation in large empirical datasets. We show that when using an appropriate transform on the branch lengths, so that the likelihood function is well approximated at the tails, the approximate method produces reliable estimates allowing Bayesian rate model selection in datasets with hundreds of taxa.

## Simulation Analysis

### Simulation strategy

#### Overview

We use the bifurcating, rooted phylogeny of Fig. 1, with *s* = 9 species and *n* = 2*s* − 2 = 16 branches, to simulate molecular alignments under the strict (STR), independent log-normal (ILN) and geometric Brownian motion (GBM) rate models. Let *r*_*i*_ be the molecular evolutionary rate (in units of substitutions per site per time unit) and Δ*t*_*i*_ be the time duration of the *i*-th branch. Simulating under a relaxed-clock model requires sampling *n* rates from the appropriate statistical distribution and then calculating the length of each branch in substitutions per site: *b*_*i*_ = *r*_*i*_Δ*t*_*i*_. The resulting phylogeny, with branch lengths in substitutions per site, is then used to simulate the molecular sequence alignments with the Evolver program from PAML (Yang 2007). Each simulated alignment is then analysed with the computer program MCMCTree (Yang 2007) for MCMC Bayesian inference of node ages and branch rates under each rate model. Marginal likelihoods and posterior probabilities for each rate model under each simulated alignment are then estimated using the stepping-stone method (Xie et al. 2011).

**Figure 1.**
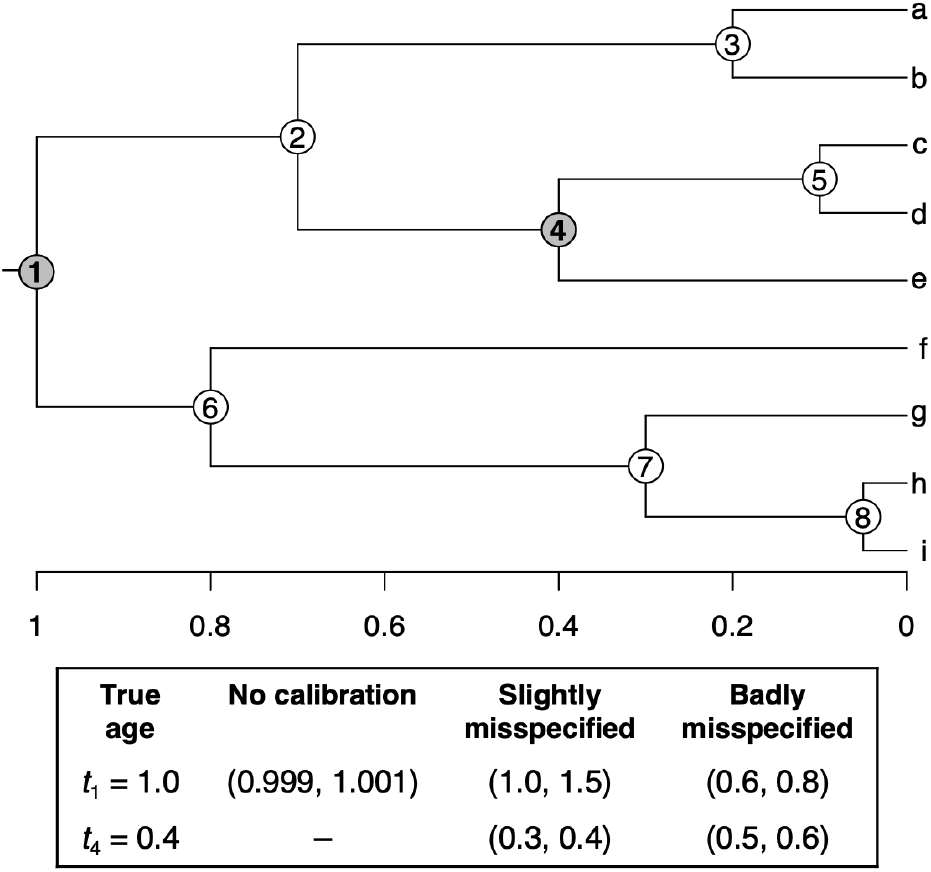
Phylogeny used to simulate nucleotide alignments under rate models. The phylogeny has eight internal nodes, 1, …, 8, with true ages *t*_*i*_ = 1, 0.7, 0.2, 0.4, 0.1, 0.8, 0.3, and 0.05. Bayesian rate model selection analyses are carried out using three fossil calibration set-ups: (1) No calibrations, i.e., the root age, *t*_1_, is fixed to one, which, in MCMCtree, is done by using a narrow uniform distribution between 0.999 and 1.001; (2) slightly misspecified calibrations, in which the true ages, *t*_1_ and *t*_4_, are at the bounds of the corresponding calibration densities; or (3) badly misspecified calibrations, in which the true ages, *t*_1_ and *t*_4_, are not contained within the corresponding calibration densities. The tree topology and node ages are the same used by Yang and Rannala (2006) in their simulation analyses.

#### Simulating rates

We have developed a new R package, simclock, to simulate rates on a phylogeny under the three rate models. In the ILN model, we sample the *n* independent, identically distributed rates from the joint log-normal distribution

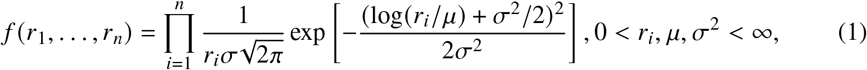

where *μ* is the mean rate and *σ*^2^ is the log-variance, which controls the departure from the strict clock. When *σ*^2^ is close to zero, the branch rates are close to *μ*, whereas when *σ*^2^ is large, there will be large rate variation among branches and the molecular clock will be seriously violated.

To simulate rates under the GBM model, we use the GBM density specification by Rannala and Yang (2007). Let *i, j* be two sister branches and *a* their parent branch, let *y*_*i*_ = log *r*_*i*_ and **y** = (*y*_*i*_, *y* _*j*_). Then *y*_*i*_ and *y* _*j*_ have a joint normal distribution conditioned on *y*_*a*_ (i.e., *r*_*i*_ and *r* _*j*_ have a joint log-normal distribution conditioned on *r*_*a*_), with density

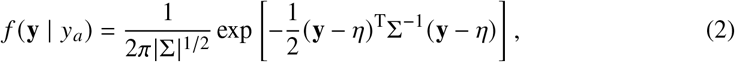

where

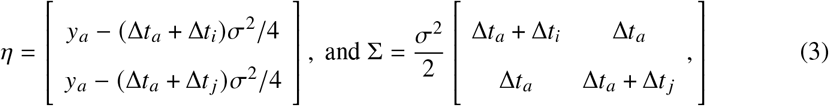

are the mean vector and covariance matrix respectively. Note that *T*_*i*_ = (Δ*t*_*a*_ + Δ*t*_*i*_)/2 is the time span between the midpoint of the parent branch *a* and the midpoint of its daughter branch *i* (see Fig. 1 in Rannala and Yang 2007). Thus, *σ*^2^*T*_*i*_ is the amount of accumulated variance for the log-rate of the daughter branch. When *σ*^2^*T*_*i*_ is close to zero, the daughter rate is close to the parent rate, whereas when *σ*^2^*T*_*i*_ is large, the daughter rate diffuses away from that of the parent, and the molecular clock is seriously violated. Note the two branches around the phylogeny’s root do not have a parent branch. In this case we set *y*_*a*_ = log *μ* and Δ*t*_*a*_ = 0. Thus, *μ* is the expected rate for the phylogeny. To sample the *n* rates, we start at the root and sample rates for its two daughter branches, *r*_*i*_ = exp *y*_*i*_ and *r* _*j*_ = exp *y* _*j*_, using Eq. (2). We then traverse the remaining nodes in the phylogeny and sample daughter rates after their parent rates have been sampled, resulting in the *n* required rates.

For the STR model, *r*_*i*_ = *μ*, for all branches. Note the STR model is a special case of both ILN and GBM when *σ*^2^ = 0.

#### Simulating alignments

In relaxed-clock dating, when both the number of sites and the number of loci (i.e., the number of alignment partitions) analysed approach infinity, the posterior of divergence times converges to a limiting distribution in which the uncertainties of time estimates are proportional to their posterior means (Rannala and Yang 2007). Rannala and Yang (2007) show that five loci already show good convergence to the limiting distribution, with 30 loci leading to excellent convergence (Fig. 4 in their paper). Thus, to study the asymptotic properties of Bayesian rate model selection, we simulate alignments with *L* = 1, 2, and 5 loci, with each locus having 1,000 sites. This set up is the largest we can analyse in reasonable time given expensive MCMC calculation of marginal model likelihoods but with reasonable convergence to the limiting distribution.

Let *A* be an alignment to be simulated, partitioned into *L* loci. Each locus evolves independently, with its own set of rates for branches. For each locus, we sample a value for the locus rate, *μ*_*h*_, *h* = 1, …, *L*, from the gamma distribution Γ(*α* = 2/*L, β* = 2/*L*), which has mean *α*/*β* = 1 and variance *α*/*β*^2^ = *L*/2. This dependency of the gamma density on *L* is used for compatibility with the gamma-Dirichlet prior used in MCMCtree (dos Reis et al. 2014). Next, for each locus, we sample a value for the log-rate variance parameter, 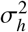, from the gamma distribution Γ(2/*L*, 20/*L*), with mean 0.1 and variance *L*/200. This sampling density leads to values of 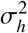 that are small and close to 0.1, thus the resulting simulated rates are similar to each other, and the phylogeny is clock-like (CL). We label simulations using this sampling density as ILN-CL or GBM-CL. Alternatively, we may also sample 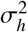 from the gamma density Γ(2/*L*, 2/*L*) with mean 1 and variance *L*/2. This leads, in many simulations, to values of 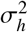 that are very large, with simulated rates being very different to each other, resulting in phylogenies in which the clock is seriously violated (SV). We label simulations using this sampling density as ILN-SV or GBM-SV. Once the values of *μ* = (*μ*_1_, …, *μ*_*L*_) and 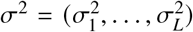 have been set, we use function `simclock::relaxed.tree` to simulate branch rates for each locus. The resulting locus trees, with branches in substitutions per site, are then used with the Evolver program from PAML (Yang 2007) to simulate *L* loci with 1,000 sites each under the Jukes and Cantor (1969) substitution model.

In total we simulated alignments from five rate model configurations (STR, ILN-CL, ILN-SV, GBM-CL, GBM-SV) and three locus configurations *L* = 1, 2, and 5, for a total of 5 × 3 = 15 simulation configurations. For each configuration, 1,000 alignments were simulated, resulting in 15,000 simulated alignments.

#### Bayesian rate model selection and rate prior

Let *M* be a rate model (one of STR, ILN or GBM). The posterior of divergence times (node ages), given the alignment, *A* = (*A*_1_, …, *A*_*L*_), and rate model, is

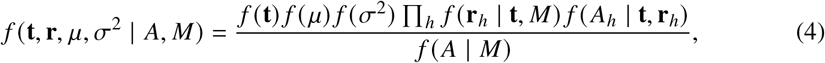

where *f* (**t**) is the prior on times, *f* (*μ*) *f* (*σ*^2^) Π_*h*_ *f* (**r**_*h*_ | **t**, *M*) is the prior on the rates and rate model parameters, *f* (*A*_*h*_ |**t, r**_*h*_) is the likelihood for the *h*-th locus in the sequence alignment, an

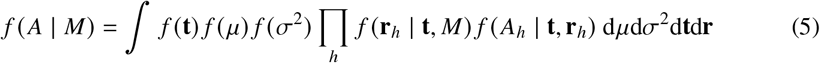

is the marginal-likelihood of the data, which depends on the rate model used to construct the rate prior. Selecting the best rate model for a given data set *A*, requires computing its marginal likelihood three times, once for each rate model. The posterior model probability is then

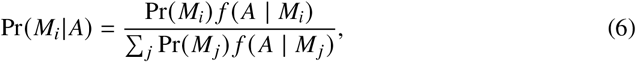

where Pr(*M*_*i*_) is the prior model probability. Here we use Pr(*M*_*i*_) = 1/3. We make a few observations about construction of the rate prior:

1. For the STR model, *f* (**r**_*h*_ | **t**, *M*) = 1 and *f* (*σ*^2^ 177 | *M*) = 1.
2. For the ILN model, *f* (**r**_*h*_ | **t**) = *f* (**r**_*h*_) is calculated using the joint log-normal density of Eq. (1).
3. For the GBM model, *f* (**r**_*h*_ | **t**) is calculated using the log-normal density associated with Eq. (2),

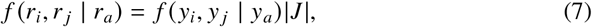

where |*J* | is the Jacobian of the (*r*_*i*_, *r* _*j*_) → (*y*_*i*_, *y* _*j*_) transform (Rannala and Yang 2007).

The joint prior on all rates is then

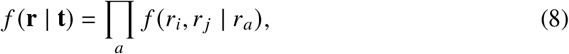

where the product is over the *s* − 1 internal nodes in the phylogeny.

4. The joint prior *f* (*μ*) *f* (*σ*^2^) over loci is calculated using the gamma-Dirichlet density (dos Reis et al. 2014). This prior uses a gamma density on the mean across locus rates, 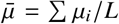, and on the mean log variance 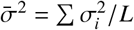, and then uses a Dirichlet distribution to partition the uncertainty across the loci. For the locus rates, we use 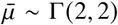 with *α* = 1 in the Dirichlet. For the log-variance, we use 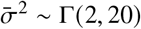, for clock-like datasets, or 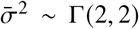, for the severely violated clock datasets. In both cases *α* = 1 in the Dirichlet is used.

#### Fossil calibrations

Calculation of marginal likelihoods and posterior rate model probabilities are carried out under three fossil calibration strategies (Fig. 1):

##### 1. No calibrations

The age of the root is fixed to one, *t*_1_ = 1, and no fossil calibrations are applied to any other nodes. The prior on the age of the remaining nodes is given by a birth-death process with parameters *μ* = 1, *λ* = 1, *ρ* = 0, resulting in a uniform kernel (Yang and Rannala 2006). In this case the model appears to be fully identifiable (see discussion), and we expect posterior model probabilities to converge to one for the true model as the number of loci increases.

##### 2. Slightly misspecified calibrations

The root is calibrated with the uniform distribution (1.0, 1.5), which has mean 1.25, thus this calibration suggests a root age that is older than the true age, *t*_1_ = 1. On the other hand, node 4 has the uniform calibration (0.3, 0.4), which has mean 0.35 which is younger than the true age *t*_4_ = 0.4. Thus the two calibrations are in conflict and push the node ages in different directions, and the true node ages are barely contained within the calibration bounds. The prior on the remaining node ages is set using the same birth-death kernel as above. In this case we expect the conflict between the two fossil calibrations to adversely affect the convergence of posterior model probabilities

##### 3. Badly misspecified calibrations

The root has the uniform calibration (0.6, 0.8), which has mean 0.7 and is thus very different than the true age, *t*_1_ = 1. Node 4 has the uniform calibration (0.5, 0.6) which has mean 0.55 and is also very different to the true age, *t*_4_ = 0.4. Here the two calibrations are in big conflict and push the node ages close together. Furthermore, neither calibration contains the true node age. The prior on the remaining node ages is set as above. In this case we expect the strong conflict between the two fossil calibrations to severely affect convergence of posterior model probabilities.

In cases 2 and 3, the calibration bounds are soft, that is, there are small tail probabilities, *p*_*l*_ = *p*_*u*_ = 2.5%, extending beyond the lower and upper bounds. This means the posterior distribution of node ages can converge to values outside the calibration bounds, making the analysis robust to misspecification of fossil calibrations (Yang and Rannala 2006).

#### MCMC sampling

The marginal likelihood (Eq. 5) cannot be calculated analytically and we must use MCMC sampling methods to estimate it. However, sampling the marginal likelihood (which is the likelihood averaged over the prior) is difficult because the prior is usually diffuse (it is spread over a large area of parameter space) while the likelihood is usually concentrated (over a small area of parameter space), and thus naive MCMC samplers will tend to obtain too many samples over the broad prior range and too few samples over the concentrated likelihood area, thus providing poor estimates with large sampling errors (Yang 2014, Ch.: 7). Here we use the stepping-stone method, which uses power posteriors to sample over the log-marginal likelihood, 𝓁 = log *f* (*A* | *M*) (Xie et al. 2011). The power posterior is given by *q*_*β*_ = *f* (**t**) *f* (*μ*) *f* (**r** | **t**, *M*) *f* (*σ*^2^ | *M*) *f* (*A* | **t, r**) ^*β*^ with 0 ≤ *β* ≤ 1. When *β* = 0, the power posterior reduces to the prior, and when *β* = 1 to the posterior. Thus, by selecting *n* values of *β* between 0 and 1, one can sample likelihood values (using MCMC) from the power posteriors in a path from the prior to the posterior. The sampled likelihoods are then used to estimate 𝓁 (see Xie et al. 2011 for details). Here we use *n* = 8 values of *β*, which requires running 8 independent MCMC samples to calculate one marginal likelihood value. We use MCMCTree to obtain the required MCMC samples, and we use the `mcmc3r` R package (dos Reis et al. 2018) to choose the *β* values and to calculate 𝓁 and Pr(*M* | *A*). The likelihood, *f* (*A* | **t, r**), is calculated exactly under the Jukes and Cantor (1969) model in all cases. We summarise the results of each simulation configuration by calculating the median (50% quantile) and interquartile range (25% and 75% quantiles) of Pr(*M* | *A*) among the 1,000 simulation replicates. For example, a median of 0.9 means in 50% of simulations the correct rate model was selected with Pr(*M* | *A*) ≥ 0.9. On the other hand, a 25%-quartile of 0.8 means in 75% of simulations the correct model had Pr(*M* | *A*) ≥ 0.8 (or Pr(*M* | *A*) ≤ 0.8 in the remaining 25%).

We note our analysis strategy required a very large number of MCMC chains: Each one of our 15,000 simulated alignments was analysed under each of 3 rate models, each rate model under three calibration strategies, with each combination requiring 8 *β* values. This gives a total of 15, 000 × 3 × 3 × 8 = 1.08 million MCMC chains. These chains were run in parallel over several months at QMUL’s Apocrita high-performance computing cluster. Configuration of the MCMC sampler in MCMCtree (step size, number of generations, and sampling frequency) was determined in test runs of the program. Convergence of the marginal likelihood estimates was assessed by plotting the mean log-likelihoods sampled under each *β* value against the *β* value, and by calculating the standard error of the marginal likelihood estimate using the stationary bootstrap (Álvarez-Carretero et al. 2022). A sample with good convergence will have small standard errors and the points in the plot will fall in a smooth curve from the prior (*β* = 0) towards the posterior (*β* = 1).

### Simulation results

Figure 2 summarises the results from the simulation analysis. When the age of the root is fixed to one, *t*_1_ = 1, and when no fossil calibrations are used, posterior model probabilities for the correct rate model converge to one in all cases (white dots, Fig. 2). When fossil calibrations are used, and when these are slightly misspecified, rate model selection is robust to the fossil misspecification and posterior model probabilities still converge to one for the correct model (black dots, Fig. 2). However, in this case, the rate of convergence is slightly worse when the true model is either STR (Fig. 2A-A’) or GBM (Fig. 2C-C’), but not for the ILN model (Fig. 2B-B’). On the other hand, when fossil calibrations are badly misspecified, posterior model probabilities converge to wrong values in all cases (asterisks, Fig. 2), with the worst behaviour for the STR model, for which model probabilities converge to zero for five loci even despite this being the correct model. The bad calibrations case is very pathological, as in many cases, a wrong model will have a higher posterior probability than the correct model, even when analysing five loci. Because in real data analyses it is not possible to guarantee that all fossil calibrations are correct, we suggest workers select the rate model in an analysis without fossil calibrations, as our simulations show this analysis setup shows the best behaviour in all cases (Fig. 2).

**Figure 2.**
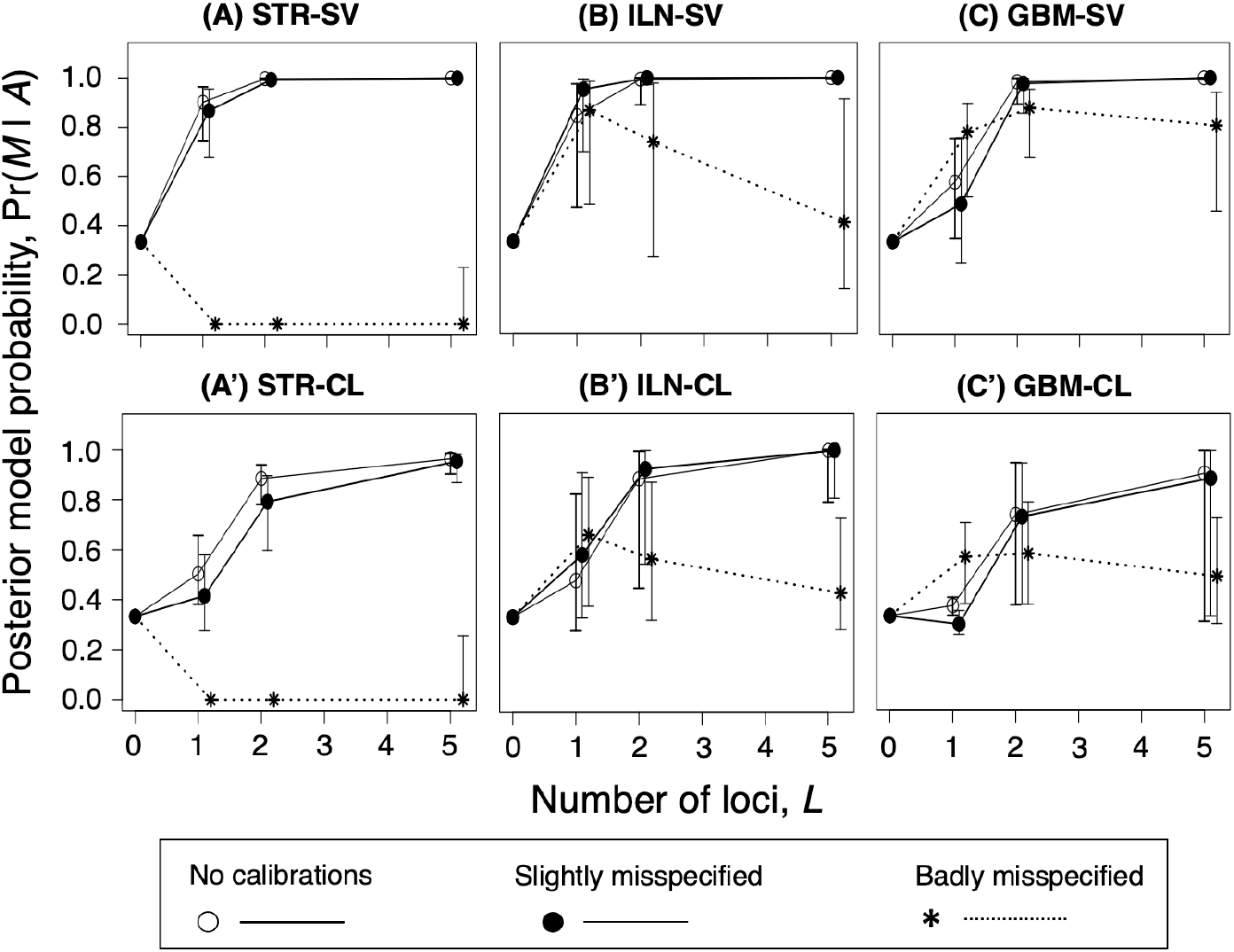
Posterior rate model probabilities for the true model in simulated datasets. The true models are (A, A’) STR, (B, B’) ILN, and (C, C’) GBM. Datasets are analysed under three fossil calibration strategies (no calibration, slightly misspecified, and badly misspecified calibrations). Each dot is the median of Pr *M|A* for the true model across 1,000 simulated datasets, and the vertical bars are the corresponding interquartile range across the simulations. Note that in (A) and (A’), the simulated datasets are the same, but the prior on 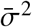 has either a high variance (seriously violated density, SV) or low variance (clocklike density, CL) when the data are analysed under the ILN and GBM models. For *L* = 0, the dots are plotted at the prior model probability, Pr(*M*) = 1/3. Likelihood is calculated exactly in all cases.

Datasets analysed using the clock-like prior, 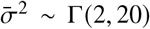, showed slower convergence in posterior probability for the correct model (Fig. 2A’-C’), when compared to the seriously violated clock prior, 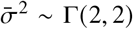 (Fig. 2A-C). The reasons for this are not clear, but we note the clock-like prior is more informative, i.e., it has a smaller variance, than the seriously violated clock prior. Thus, our results here suggest a diffuse prior on 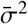 should be preferred.

Figures 3 and 4 show posterior model probabilities for the true model vs. true mean locus rate, 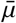, and true mean rate parameter, 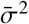, for each simulated dataset. The patterns in the figure are very instructive. For example, it is difficult to ascertain the right model when the true mean rate is very high, and in particular when the number of loci is low. Furthermore, among the clock-like simulations, it is hard to distinguish the right model when the true model is GBM, and this effect is reduced as the number of loci is increased. We emphasise each dataset and locus are simulated with specific values of *μ* and *σ*^2^ sampled from the appropriate simulation densities, and their posterior estimates thus reflect the underlaying distribution. For computational reasons we could not test larger numbers of loci (i.e., *L* > 5) but it is clear from the asymptotic analysis (2) that the probability of the correct model should converge to one with more than five loci when no fossil calibrations are used.

**Figure 3.**
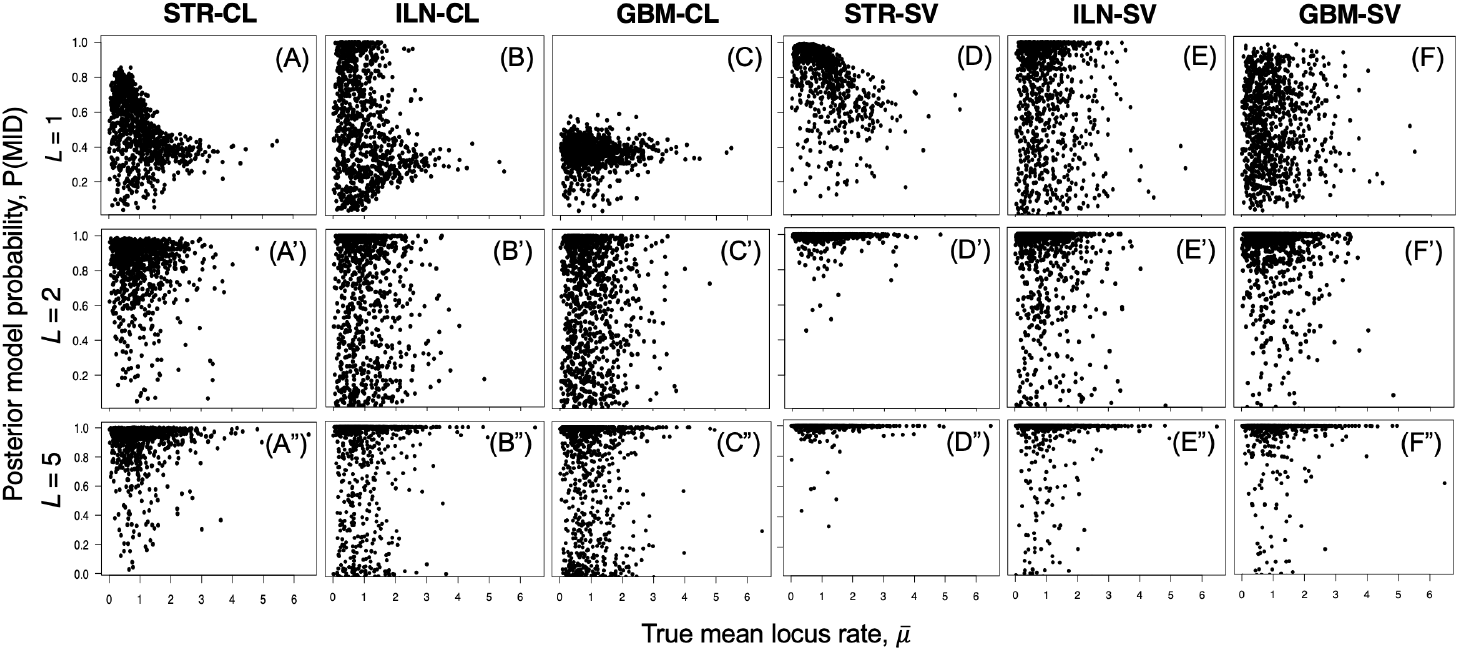
Posterior rate model probabilities for the true model vs the true mean locus rate. Each dot represents the posterior probability of the true model for a single simulated dataset (i.e., each panel contains 1,000 dots corresponding to 1,000 simulations for the particular parameter setting), with data simulated under the clock-like (CL) and seriously violated (SV) rate densities. The data are analysed without fossil calibrations (i.e. the root age is fixed to one).

**Figure 4.**
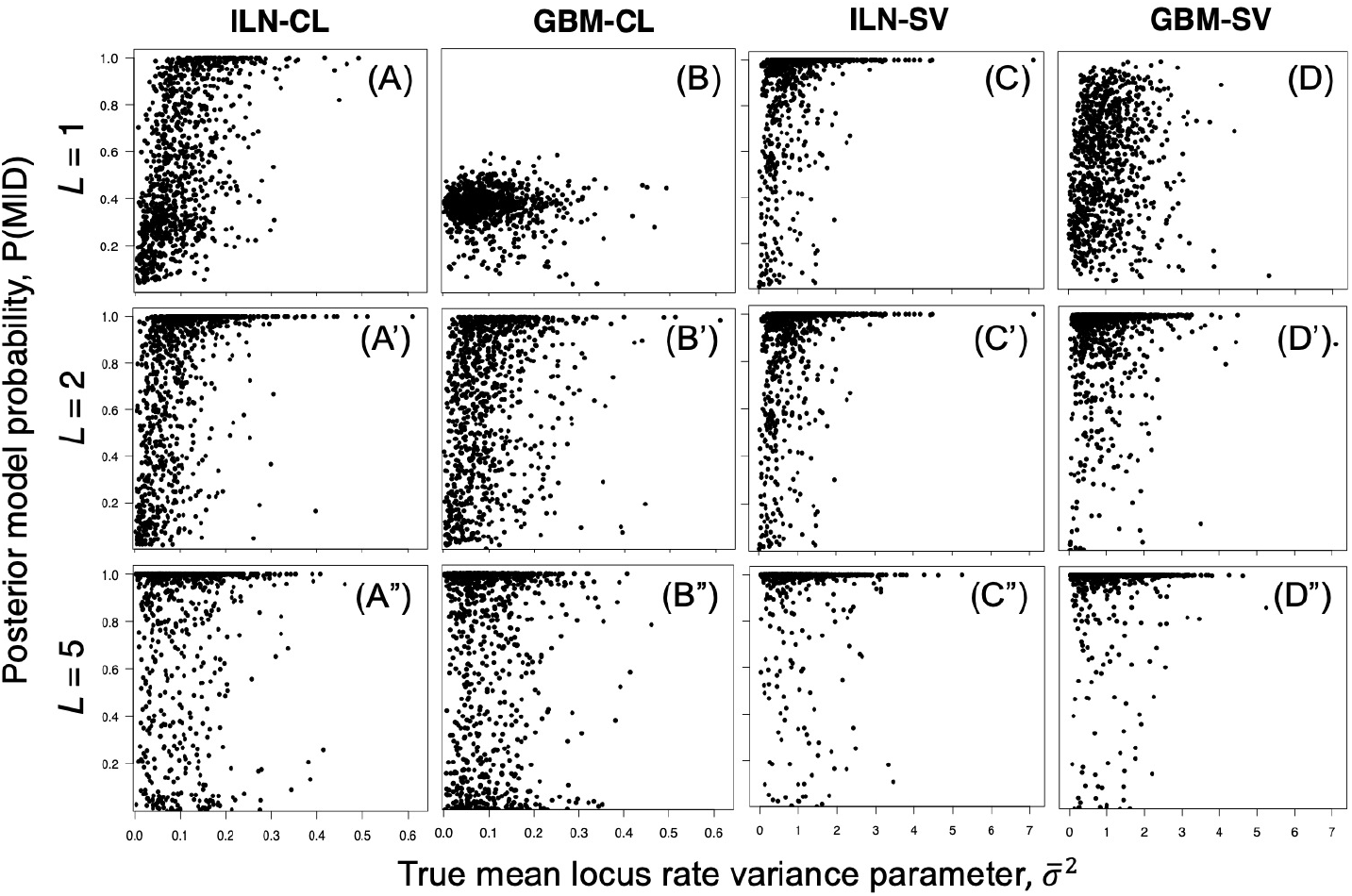
Posterior rate model probabilities for the true model vs the true mean drift parameter. See legend of Fig. 3.

## Approximations for Fast Likelihood Computation on Large Phylogenies

Thorne et al. (1998) (see also Seo et al. 2004) suggest approximating the likelihood to speed up computation during MCMC sampling of divergence times, with dos Reis and Yang (2011) suggesting transformation of branch lengths to improve the accuracy of the approximation. The approximation can provide speed-ups of up to 1,000x compared with exact likelihood calculation depending on the dataset (Battistuzzi et al. 2011). However, in the original approximation by Thorne et al. (1998), the approximation error can be substantial at the tails of the likelihood function, with the error worsening as the distance away from the likelihood peak increases (dos Reis and Yang 2011: Fig. 1). The transforms proposed by dos Reis and Yang (2011) reduce the approximation error at the tails, with the Arcsine-based transform providing the smallest errors. When conducting MCMC sampling of divergence times under the posterior (Eq. 4), the errors in the likelihood tail appear unimportant because values at the tail are rarely sampled (dos Reis and Yang 2011), and posterior estimates from the approximate method are indistinguishable from the exact method in large datasets (dos Reis and Yang 2011). On the other hand, tail errors are expected to be important in marginal-likelihood calculation. For example, in the stepping-stone method, tail values will be sampled often when *β* is close to zero in the power posterior. It is not clear whether tail errors under the branch-length transforms are small enough to obtain reliable estimates of the marginal likelihoods. Furthermore, when the clock is seriously violated, and when the data are analysed under the clock, the approximate method does not work well because sampled branch-lengths are very far from the likelihood peak (dos Reis and Yang 2011). Here we examine the accuracy of the approximate method when estimating marginal likelihoods for rate models and use the approximate method to select for the best rate model in phylogenies with several hundred taxa.

### Accuracy of marginal likelihood estimation under the approximate method

We use two real datasets. The first is the 12 mitochondrial protein-coding genes from seven ape species analysed by Yang and Rannala (2006). The genes are divided into three partitions according to codon position, with each partition having 3,331 sites. The second dataset is the 1st and 2nd codon positions from 10 nuclear protein-coding genes from 72 mammal species. This is a subsample of 10 genes out of the 15,268 analysed by Álvarez-Carretero et al. (2022), with the number of sites ranging from 406 to 4,946 among the 10 genes. Thus, the two datasets result in 13 partitions: three partitions for the ape data, and ten partitions for the mammal data. For each partition, we calculate marginal likelihoods for each rate model, both under the exact and approximate likelihood methods, resulting in 13 × 3 × 2 = 78 marginal likelihood calculations. All 13 partitions are analysed under the HKY+Γ model (Hasegawa et al. 1985; Yang 1994). We note that in the exact method, the *κ* and *α* parameters from the HKY+Γ model are sampled during MCMC, while they are fixed at their maximum likelihood estimates during MCMC sampling with the approximate method (dos Reis and Yang, 2011). This means the marginal likelihood estimates between the exact and approximate approaches cannot be compared directly: the approximate method will result in larger marginal likelihoods because in this case integration over the priors on *κ* and *α* does not take place. Instead, we compare the log-Bayes factors between models. Let 𝓁_*i*_ and 𝓁_*j*_ be the log-marginal likelihoods for two models. The log-Bayes factor, or marginal log-likelihood difference, is Δ𝓁_*i j*_ = 𝓁_*j*_ − 𝓁_*i*_.

Figure 5 shows the log-Bayes factors calculated for the 13 partitions and 3 rate models under the exact and approximate approaches. In all 13 partitions, the molecular clock is seriously violated, and the approximate Δ𝓁 values calculated in this case are not reliable. For example, the largest discrepancies in Δ𝓁 values can be as large as 100 log-units (Fig. 5A-D). On the other hand, when comparing the two relaxed-clock models (ILN vs GBM), the approximations using branch transformations (ARCSINE, SQRT and LOG) perform very well, showing only slight errors in the estimates of Δ𝓁 (Fig. 5A’-C’). The approximation without branch-transformation (NT) does show substantial errors in the calculation of Δ𝓁 in two comparisons (Fig. 5D’). These results are re-assuring as they demonstrate that approximating the branch lengths under the transform are good enough to approximate Δ𝓁 in real datasets. As is evident from Figure 5, the ARCSINE transform provides the best approximation, so we suggest this approximation is used to calculate marginal likelihoods for relaxed-clock models in very large datasets. We do not recommend using the NT method for any comparisons, or any of the approximate methods if comparisons with the strict clock are required.

**Figure 5.**
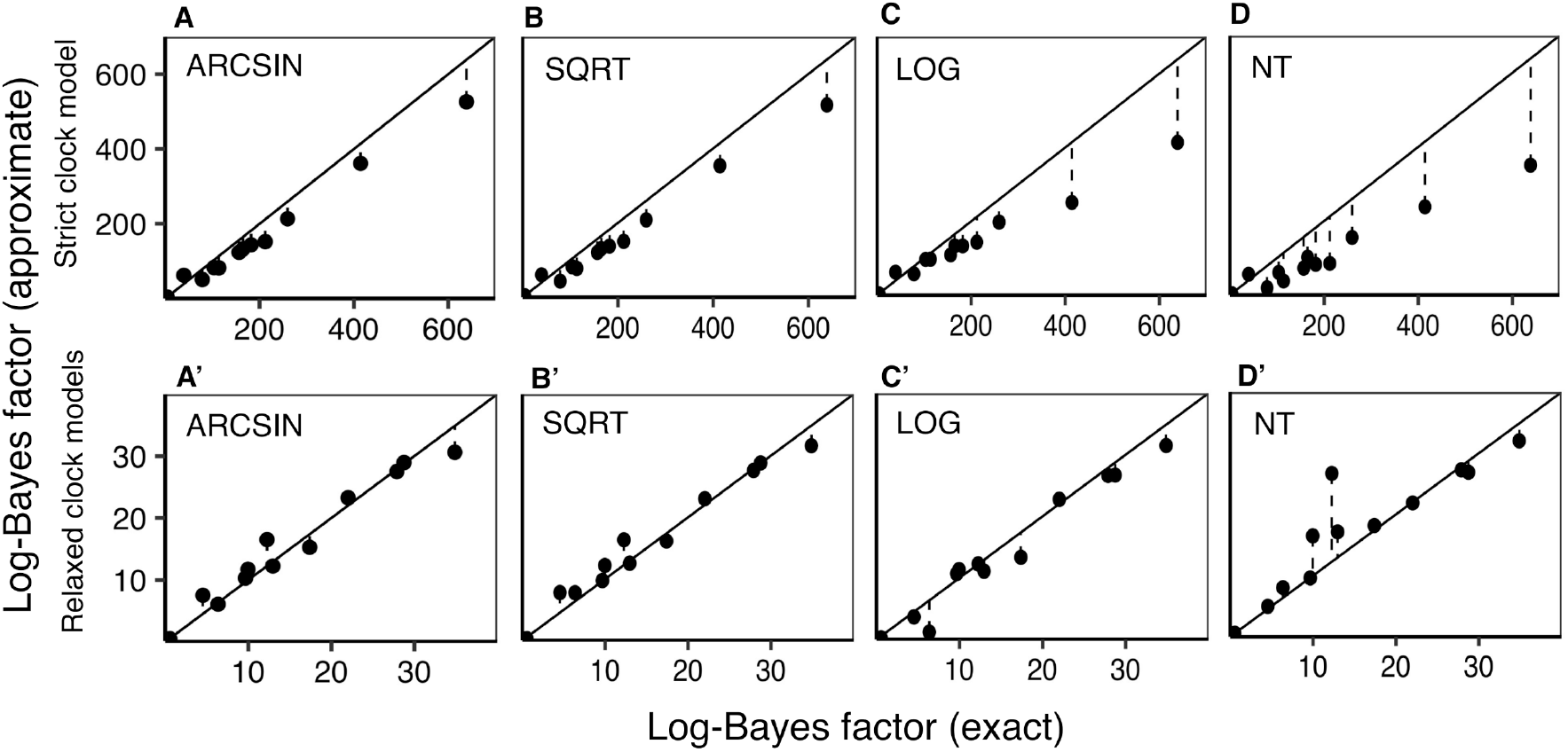
Comparison of log-Bayes factors (Δ𝓁 of rate models under exact and approximate likelihood calculation. The data are alignments of ten nuclear genes from 72 mammal species and the three codon positions from the mitochondrial proteins from seven ape species (13 partitions in total, analysed separately). (A-D) Comparisons between the GBM and STR rate models for four methods to transform branch lengths (ARCSIN, SQRT, LOG and NT, see dos Reis and Yang 2011 for details). (A’-D’) Comparisons between the GBM and ILN models for the four transform methods. Each dot is the Δ𝓁 value from for a given model comparison on a single alignment. Notice the expanded range of values for the strict-clock analysis (top row) in the x-and y-axes. This is due to the poor performance of the STR model when compared to the GBM model in this dataset.

### Application of the approximation to select the rate model in large datasets

dos Reis et al. (2018) and Barba-Montoya et al. (2018) used marginal likelihood to select the best rate model in Primate and flowering plant phylogenies respectively. Because those authors estimated the likelihood exactly, they were restricted to small datasets in their Bayesian rate model selection analyses: 10 species in dos Reis et al. (2018) and 10 species in Barba-Montoya et al. (2018), a dramatic subsampling of their large phylogenies with 372 and 644 species respectively. Thus, an open question remains, would have those authors selected the same rate model if all species had been analysed? Here we re-analysed the two phylogenies to select for the best rate model using all species and the approximate method. Table 1 shows the results. In the original analysis by dos Reis et al. (2018) the preferred rate model was GBM for the 10 species across five out of six alignment partitions, with the sixth partition, mitochondrial 3rd codon positions, supporting the STR model. This result is confirmed here for all 372 species, except for the 3rd codon positions, which support the GBM model in the full dataset (Table 1). Barba-Montoya et al. (2018) found best support for the ILN model among 10 species and across three alignment partitions. This result is confirmed here for all 644 species and all partitions (Table 1). Both large datasets appear very informative about the rate model, as the posterior probabilities for the best model are always one in each case, as the marginal likelihood differences across models are very large (Table 1).

**Table 1:** Bayesian rate model selection in large phylogenies using approximate likelihood

| Dataset<sup>a</sup> | Partition<sup>b</sup> | $\ell_{\text{STR}} \pm \text{SE}^c$ | $\ell_{\text{ILN}} \pm \text{SE}^c$ | $\ell_{\text{GBM}} \pm \text{SE}^c$ |
| --- | --- | --- | --- | --- |
| Primates<br>(372 species) | Mit 1+2 | $-1249.87 \pm 0.90$ | $-980.28 \pm 1.03$ | **$-938.84 \pm 0.89$** |
| | Mit 3 | $-3240.69 \pm 15.98$ | $-1303.65 \pm 0.36$ | **$-1187.9 \pm 0.64$** |
| | Mit RNA | $-795.94 \pm 0.16$ | $-660.44 \pm 0.37$ | **$-638.33 \pm 0.62$** |
| | Nuclear 1+2 | $-1123.02 \pm 0.57$ | $-788.19 \pm 0.57$ | **$-730.76 \pm 0.70$** |
| | Nuclear 3 | $-1245.53 \pm 0.81$ | $-855.93 \pm 0.63$ | **$-773.40 \pm 0.61$** |
| | Nuclear NC | $-1717.58 \pm 6.38$ | $-982.28 \pm 0.55$ | **$-892.60 \pm 0.84$** |
| Angiosperms<br>(644 species) | Plastid 1+2 | $-29877.16 \pm 31.19$ | **$-2877.76 \pm 0.72$** | $-2958.10 \pm 1.32$ |
| | Mit 1+2 | $-4667.87 \pm 2.50$ | **$-1607.79 \pm 0.28$** | $-1643.91 \pm 0.52$ |
| | Nuclear RNA | $-6838.14 \pm 1.98$ | **$-1697.54 \pm 0.25$** | $-1763.13 \pm 0.42$ |

a. The Primates data is from dos Reis et al. (2018) and the Angiosperm (flowering plants) data from Barba-Montoya et al. (2018). The number of species indicated is the total number in the phylogeny, but the actual number in each partition may differ (see original publications for details). b. 1+2: 1st and 2nd codon positions, 3: 3rd codon positions. NC: non-coding. c. The ARCSIN transform and root calibration (0.999, 1.001) are used in all cases. The standard error (SE) for the log-marginal likelihood estimates is calculated using the stationary bootstrap (Politis and Romano 1994; Álvarez-Carretero et al. 2022). The model with the largest marginal likelihood is shown in boldtype face. The posterior probability for the best model is 1 in all cases, as the marginal likelihood differences are very large.

Selecting the most appropriate rate model in these datasets appears important because the two rate models result in substantially different evolutionary timelines. For example, in the case of Primates, dos Reis et al. (2018) inferred an age for the crown group of 74.4 Ma (95% Credibility interval: 79.2–70.0 Ma) using the GBM model, while this was substantially older at 90.6 Ma (95% CI: 98.1–83.7 Ma) under the ILN model. Similar discrepancies in time estimates for the two rate models were seen by Barba-Montoya et al. (2018) in their flowering plant analyses. Finally, we note errors in marginal likelihood estimates due to MCMC sampling can be noticeable. For example, for the Primate mitochondrial 3rd codon position partition under the STR model, the estimated standard error (SE) of 𝓁 was 15.98 log-units (Table 1). In this case, the error is unimportant as the log-likelihood difference between models is much larger (over 2,000 log-units between STR and GBM). However, an error of 1 log-unit would be too large if, for example, the true posterior probability for the best model is around 90%. As a conservative criterion, we recommend the sum of the SEs be much less than the absolute value of the difference in log-marginal likelihoods, i.e., SE_*i*_ + SE_*j*_ ≪ |𝓁_*i*_ − 𝓁_*j*_ |. If this is not the case, the SEs can be reduced by increasing the length of the MCMC and/or increasing the number of *β* points in the power posterior. A fourfold increase in the length of the MCMC decreases the SE by half.

## Discussion

The posterior probability of the true model is guaranteed to converge to one as the amount of data goes to infinity, if the true model is among the models tested and if certain regularity conditions are met, such as data independence and parameter identifiability (e.g., Yang and Zhu 2018). In Bayesian molecular clock dating with fossil calibrations, times and rates are non-identifiable and thus the conditions that guarantee convergence to the true model are not met. However, identifiability problems may be solved by finding functions of parameters (e.g., reparameterisations) that are identifiable (Rannala 2002). In our case, by fixing the age of the root to one, we aim to make the rates and node ages identifiable, because we are reducing the degrees of freedom of the inference problem and thus we estimate relative instead of absolute node ages. We note it appears no formal proof of identifiability of relative times and rates under relaxed-clock models exists, but the evidence from the strict clock model and MCMC simulations with infinite sites in relaxed-clock models indicate this should be the case (Rannala and Yang 2007; Yang and Rannala 2006). For the strict clock, Yang and Rannala (2006) show that when the number of sites goes to infinity, the posterior distribution of times converges to a one dimensional density and the uncertainty of node ages is proportional to their posterior means (Eq. 21 in their paper). The corollary of Yang and Rannala (2006)’s proof is that if the age of the root is fixed to one, then the density must collapse to a point estimate for the infinite data, i.e., the relative node ages are fully identifiable in this case. Under relaxed clocks, the limiting density for an infinite number of loci and infinite number of sites is not available analytically, and thus Rannala and Yang (2007) used MCMC simulation to study the asymptotic behaviour of the distribution. Their results demonstrate the distribution of node ages also approaches a limiting distribution. Thus, we suspect estimation of relative node ages is also fully identifiable under relaxed clocks, but a formal proof is still required.

Our simulations show that, when we don’t use fossil calibrations and estimate relative ages, posterior rate model probabilities asymptotically converge to one for the correct model as expected by theory. Thus, we recommend phylogeneticists interested in selecting the best rate model for their datasets use this approach. We note we did not perform simulations with good fossil calibrations. By good fossil calibrations we mean those in which the true node age is well contained within the calibration densities, for example, when using conservative maximum bounds that aim to encompass the likeliest ages of nodes (Donoghue and Benton, 2007). It is clear this case would be well-behaved, rate estimates for branches would converge to their true values, and posterior probabilities would converge to one for the correct model as for the no-fossil calibrations case. However, in real data analyses, it appears difficult to be sure about whether all calibrations are good or not. In our experience, conflicts between calibrations and posterior estimates are common (e.g., Fig. 3 in dos Reis et al., 2012). As we show here, if the calibrations are slightly misspecified, posterior model probabilities are still well behaved, but if they if the calibrations are badly misspecified, posterior model probabilities may be seriously compromised and the wrong model may be selected. Rate model selection with fossil calibrations may still be a worthwhile effort if they are accompanied by analyses with no calibrations: If discrepancies between the two model selection procedures arise, this may be indicative of problems with the calibrations, and further investigation on the accuracy of the calibrations may be warranted.

A limiting factor in the application of Bayesian rate model selection to real datasets has been the extreme computational expense of calculating the likelihood for large phylogenies during MCMC sampling across power posteriors. This means that relatively few studies have calculated posterior probabilities for rate models, and in all cases the datasets analysed have been small, with few species and/or few sites (Linder et al. 2011; Lepage et al. 2007; Baele et al. 2012; dos Reis et al. 2018; Barba-Montoya et al. 2018; McGowen et al. 2020; Álvarez-Carretero et al. 2022). Of course, if estimated divergence times are similar among rate models (e.g. Jarvis et al. 2014; Foster et al. 2017), then rate model selection may not be necessary. On the other hand, if large discrepancies are evident (e.g. dos Reis et al. 2018), then some sort of model selection is necessary. A computationally efficient alternative to calculating marginal likelihoods is model averaging, in which different rate models are sampled during MCMC (Li and Drummond 2012; Lartillot et al. 2016; Zhang 2022). In this case the posterior distribution of times is averaged across the rate models sampled. However, mixing of the MCMC among models may be difficult (Yang 2014: ch 7) and the posterior probability of a rate model cannot be calculated if its very low, as the model will be too rarely sampled.

In previous studies we had advised against using the approximate method for estimating marginal likelihoods due to concerns about the accuracy of the approximation in the tails of the likelihood function (dos Reis et al. 2018; Álvarez-Carretero et al. 2022). However, here we show that those concerns were too conservative, and the approximations under the ARCSINE and SQRT transform are good enough to provide reliable estimates of posterior model probabilities. Calculation speed-ups under the approximation can be quite dramatic, for example, a dataset that may require one day of MCMC sampling under the approximation, may required nearly three years under the exact method, given the 1000x speed-up reported in a benchmarking study (Battistuzzi et al. 2011). We recommend phylogeneticists interested in testing for rate models in large datasets use the approximation under the ARCSINE transform, as this provides the best approximation for the long tail of the likelihood. This may be accompanied by sanity tests using the exact method on a smaller subsample of species from the phylogeny. However, we note the approximate method must not be used if the strict clock is among the models being compared or if extremely long branches are present in the phylogeny.

Finally, we would like to point out that the approximate method may have applications beyond rate model selection. For example, Fourment et al. (2020) explored the use of similar approximations to estimate the marginal likelihood and select the best tree topology among a candidate set of trees (for example, among the set of *n* trees with highest likelihood from a maximum likelihood analysis). Thus, our results here suggest the approximation currently implemented in MCMCtree can also be used to estimate posterior model probabilities among tree topologies in very large phylogenomic datasets, datasets for which traditional MCMC sampling with tree moves would be computationally prohibitive: We may select the top *n* topologies from a maximum likelihood analysis with, say with RAxML or IQ-TREE, then use MCMCtree with the approximate method to estimate the corresponding marginal likelihoods for each tree topology and calculate their posterior probabilities. This would make Bayesian topology estimation for phylogenomic datasets with hundreds of taxa and millions of sites within reach.

## Supplementary Material

The Primate and Angiosperm datasets are available from their original data repositories at http://dx.doi.org/10.5061/dryad.c020q and https://figshare.com/s/404b70bc39656c2cf57e respectively. The scripts used in the simulation and real data analyses are available at https://github.com/M The simclock R package, to simulate relaxed rate models on phylogenies, is available from https://github.com/dosreislab/simclock. The mcmc3r R package is available from https://github.com/dosreislab/mcmc3r. This package can be used to select *β* values, and prepare MCMCtree control files, to estimate marginal likelihoods using the stepping-stone (Xie et al. 2011) and thermodynamic integration with Gaussian quadrature (Rannala and Yang 2017) methods. The package also allows calculation of the standard error of marginal likelihood estimates using analytical formula (Xie et al. 2011; dos Reis et al. 2018), and the stationary bootstrap (Álvarez-Carretero et al. 2022).

## Funding

This work was supported by Biotechnology and Biological Sciences Research Council, UK, grant BB/T01282X/1. LF was supported by Fundação de Amparo À Pesquisa do Estado de São Paulo, Brazil, grant 2018/19682-8.

## Conflicts of Interest

The authors declare no conflicts of interest.

## Notes

https://github.com/Muthubioinfo/RelaxedBF

### Competing Interest Statement

The authors have declared no competing interest.

### Summary of Updates

Text was revised to accommodate reviewer suggestions. One table and one figure were removed. The discussion section was expanded.

